# Unique cortical and subcortical activation patterns for different conspecific calls in marmosets

**DOI:** 10.1101/2024.04.09.588714

**Authors:** Azadeh Jafari, Audrey Dureux, Alessandro Zanini, Ravi S. Menon, Kyle M. Gilbert, Stefan Everling

## Abstract

The common marmoset (*Callithrix jacchus*) is known for its highly vocal nature, displaying a diverse range of different calls. Functional imaging in marmosets has shown that the processing of conspecific calls activates a brain network that includes fronto-temporal cortical and subcortical areas. It is currently unknown whether different call types activate the same or different networks. Here we show unique activation patterns for different calls. Nine adult marmosets were exposed to four common vocalizations (phee, chatter, trill, and twitter), and their brain responses were recorded using event-related fMRI at 9.4T. We found robust activations in the auditory cortices, encompassing core, belt, and parabelt regions, and in subcortical areas like the inferior colliculus, medial geniculate nucleus, and amygdala in response to these conspecific calls. Different neural activation patterns were observed among the vocalizations, suggesting vocalization-specific neural processing. Phee and twitter calls, often used over long distances, activated similar neural circuits, whereas trill and chatter, associated with closer social interactions, demonstrated a closer resemblance in their activation patterns. Our findings also indicate the involvement of the cerebellum and medial prefrontal cortex (mPFC) in distinguishing particular vocalizations from others.

**Significance Statement:** This study investigates the neural processing of vocal communications in the common marmoset (*Callithrix jacchus*), a species with a diverse vocal repertoire. Utilizing event-related fMRI at 9.4T, we demonstrate that different marmoset calls (phee, chatter, trill, and twitter) elicit distinct activation patterns in the brain, challenging the notion of a uniform neural network for all vocalizations. Each call type distinctly engages various regions within the auditory cortices and subcortical areas, reflecting the complexity and context-specific nature of primate communication. These findings offer insights into the evolutionary mechanisms of primate vocal perception and provide a foundation for understanding the origins of human speech and language processing.

## Introduction

Most primate species live in groups which provide them with socially complex structures. Vocal communication plays a key role in such groups because it allows individuals to avoid predators, interact with other group members, and promote cohesion within social groups during daily activities or travel ^1^. Vocalization in nonhuman primates (NHPs) is considered a precursor for human language, and speech perception in humans likely evolved in our ancestors by using pre-existing neural pathways that were responsible for extracting behaviorally relevant information from the vocalizations of conspecifics ^2^.

Neuroimaging studies in Old World macaque monkeys suggest that responses to conspecific calls are structured bilaterally in a gradient form along the superior temporal lobe wherein the rostral parts are predominantly activated by integrated vocalizations while the caudal parts are responsive to the acoustic features of these calls ^3^. Several studies examined the activations of neurons within the auditory cortex in response to conspecific vocalizations in macaque monkeys. Some level of selectivity for three or fewer out of seven calls was reported in all three lateral belt regions including the anterolateral (AL), mediolateral (ML), and caudolateral (CL) regions with the AL field displaying the most robust level of selectivity ^2,4^. Beyond the auditory cortex, single-unit recordings also demonstrate that the macaque ventrolateral prefrontal cortex (vlPFC) is involved in assessing acoustic features unique to conspecific vocalizations. The majority of the neurons within this area demonstrated some level of selectivity when the monkeys were presented with diverse vocalizations ^5^.

Given the challenges of studying vocal and cognitive auditory processing in general in macaques ^6,7^, New World common marmosets (*Callithrix jacchus*) have emerged as a powerful additional NHP model for vocal studies ^8^. Potentially because of their densely foliated arboreal habitat and family structure, marmosets possess a diverse array of calls that is dependent on social contexts and ecological factors ^9,10^. This rich vocal repertoire underlies their consistent engagement in acoustic communication which is also characterized by vocal turn-taking ^1^^1^, a critical feature shared with humans ^12^.

While electrophysiological recordings ^13,14^ and more recently two-photon calcium imaging^14^ have revealed a subset of neurons tuned to specific call types in auditory cortices, it is unknown whether different neural circuits are activated by different vocalizations in marmosets. If this is the case, it is possible that vocalizations produced in different contexts are processed by different brain networks. The neural substrate involved in processing long-distance calls, such as phee and twitter, would then be more similar compared to the neural circuitry predisposed to the processing of short-distance calls, such as trill, or emotionally charged ones, such as chatter. Another possibility is the involvement of different brain networks based on the acoustic features of vocalizations, with a more similar neural substrate for calls sharing more acoustic characteristics ^8^.

Recently, we developed a technique to obtain whole-brain fMRI in awake marmosets at 9.4T in response to auditory stimuli and used it to map the marmosets’ vocalization-processing network. We found that blocks of mixed conspecific vocalizations evoked stronger activations than scrambled vocalizations or non-vocal sounds in a fronto-temporal network ^15^ including auditory and cingulate cortices. Here we followed up on this approach and utilized event-related fMRI to test whether single phee, chatter, trill, or twitter calls evoked similar or distinct patterns of activation in awake marmosets. Our data show that although all calls activated a core network of cortical and subcortical areas, each call is associated with a distinct activation pattern.

## Results

We performed event-related fMRI at 9.4T in nine awake marmosets. Utilizing a “bunched slice acquisition sequence” (Fig. 1A), we collected each echo-planar imaging volume in 1.5 sec, while the effective TR was 3 sec (with 1.5 sec of silent time between volumes). We presented one of four types of marmoset vocalizations (phee, chatter, trill, or twitter) during the silent periods every 3-12 s in a pseudorandom order (Fig. 1B). Figure 1C illustrates the combined group activation in response to all four vocal stimuli compared to the baseline periods when no auditory stimuli were presented to the monkeys. The findings show robust activations in all regions of the auditory cortices including the core, belt, and parabelt at the cortical level. Subcortically, the calls activated the inferior colliculus (IC), medial geniculate nucleus (MGN), amygdala (Amy), the reticular nucleus of the thalamus (RN), and the brainstem reticular formation (RF).

**Figure 1:**
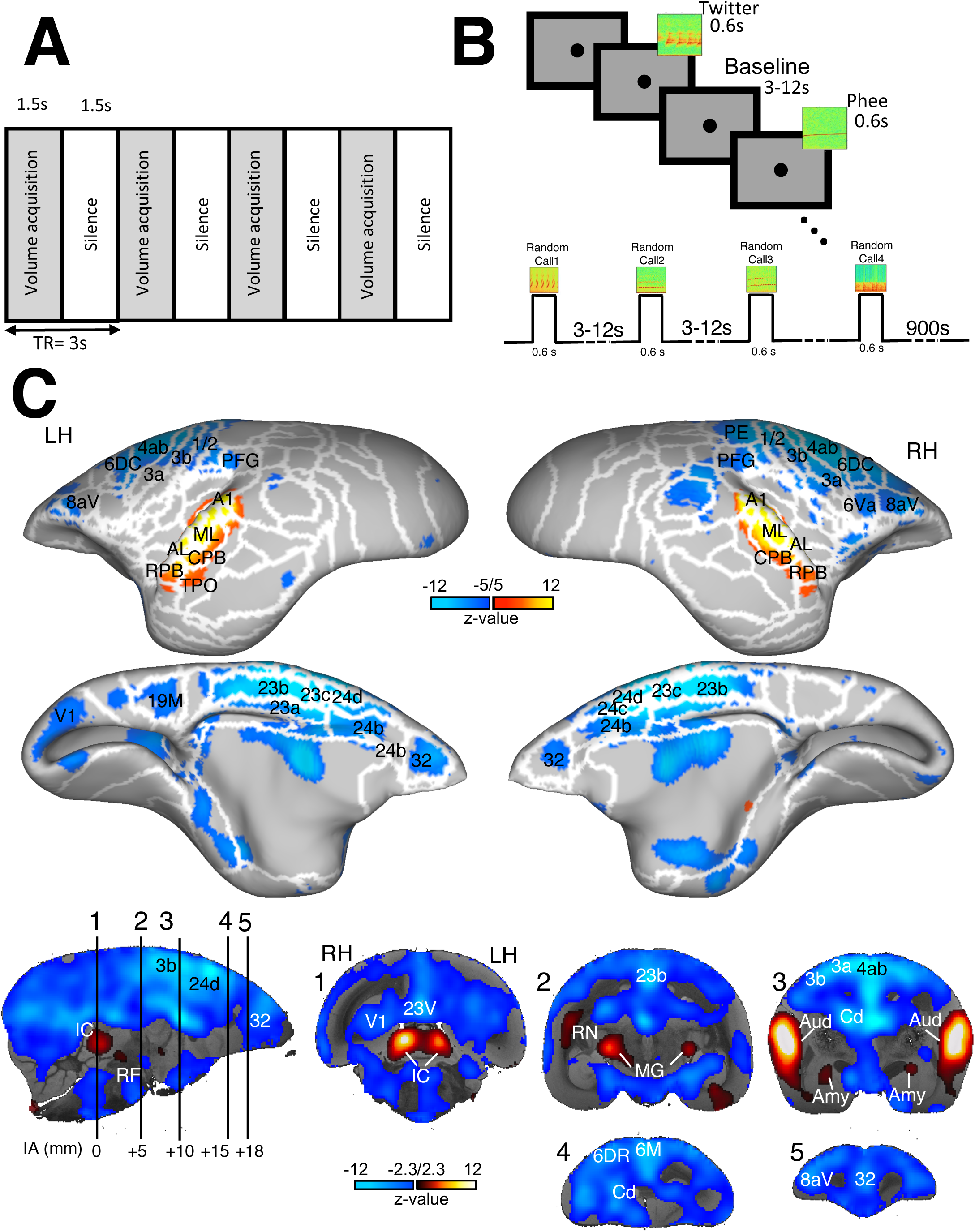
fMRI study overview. **(A)** The sequential process of a bunched slice acquisition paradigm utilized in the study. Each acquisition cycle comprises an acquisition time (*TA*) of 1.5 seconds followed by a silent period (*TS*) of equal duration, collectively constituting a repetition time (*TR*) of 3 seconds. **(B)** Graphical representation of the experimental task paradigm employed in the current study. Auditory stimuli with a duration of 0.6 seconds are randomly presented to marmoset subjects during the silent periods depicted in Fig.A with inter-stimulus intervals varying between 3 and 12 seconds, pseudorandomly chosen. **(C)** Representation of group brain activation comparison (n= 9 marmosets) for overall auditory tasks versus baseline. The upper panels depict surface maps, providing a topographical view of cortical activations. White lines delineate regions based on the atlas from Paxinos et al ^51^. The lower panels show volumetric representations at different interaural (IA) levels, overlaid onto coronal slices of anatomical MRI images. All surface maps are set to a threshold of z-scores below −5 and above 5, while volumetric maps are set to a threshold of z-scores below −2.3 and above 2.3, for deactivation and activation correspondingly. Cold color gradients indicate deactivation (negative values), while hot color gradients signify activation (positive values), representing the spatial distribution and intensity of neural responses during the auditory task. **LH**, left hemisphere; **RH**, right hemisphere; **Aud**, auditory cortex; **MG**, medial geniculate nucleus; **IC**, inferior colliculus; **Amy**, amygdala; **RN**, reticular nucleus of the thalamus; **RF**, the brainstem reticular formation; **Cd**, caudate; **A1**, primary auditory cortex area; **ML**, auditory cortex middle lateral area; **AL**, auditory cortex anterolateral area; **CPB**, caudal parabelt area; **RPB**, rostral parabelt area, **TPO**, temporo-parietal-occipital area.

Moreover, our findings in Fig. 1C show strong widespread deactivation in response to the brief vocalizations predominately in sensorimotor regions including primary motor cortex 4ab, premotor cortex areas 6DC, 6DR, 6Va, and 6M, cingulate cortices 32, 24a-d, 23a-c, somatosensory cortex areas 1/2, 3a, 3b, prefrontal area 8Av, and parietal areas PE and PFG. Additionally, Figure 2 illustrates bilateral deactivation in the cerebrocerebellum (Crll) for all vocal stimuli versus baseline.

**Figure 2:**
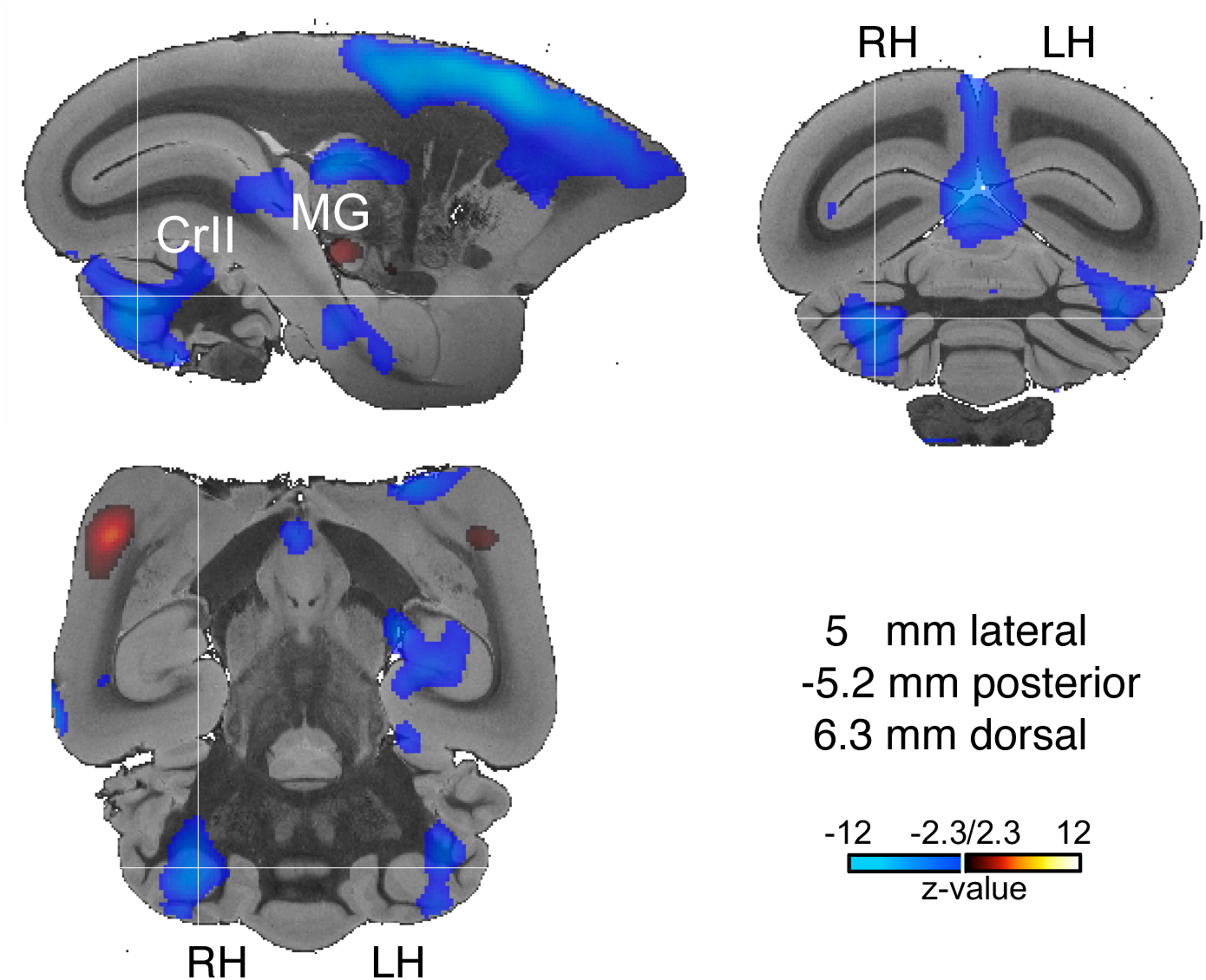
Bilateral deactivation of the cerebellum. Volumetric representation illustrating the deactivation of the cerebellum in response to the overall auditory tasks. All volumetric maps are set to a threshold of z-scores below −2.3 and above 2.3, for deactivation and activation, respectively. Cold color gradients with negative values show deactivation, while hot color gradients with positive values denote activation, representing the spatial distribution and intensity of neural responses during the auditory task. **LH**, left hemisphere; **RH**, right hemisphere; **MG**, medial geniculate nucleus; **Crll**, cerebellum.

To directly identify the activation pattern associated with a particular call, we subsequently conducted group comparisons (n = 82 runs) between the functional response for each of the four vocalizations and the baseline in Figure 3. Overall, each of the four vocalizations activated a relatively similar network in marmosets. This shared call-specific network predominantly encompassed the auditory cortices including core (primary area [A1], rostral field [R], and rostral temporal [RT]), belt (caudomedial [CM], CL, ML, AL, rostromedial [RM], rostrotemporal medial [RTM], rostrotemporal lateral [RTL]), and parabelt (rostral parabelt [RPB] and caudal parabelt [CPB]) regions.

**Figure 3.**
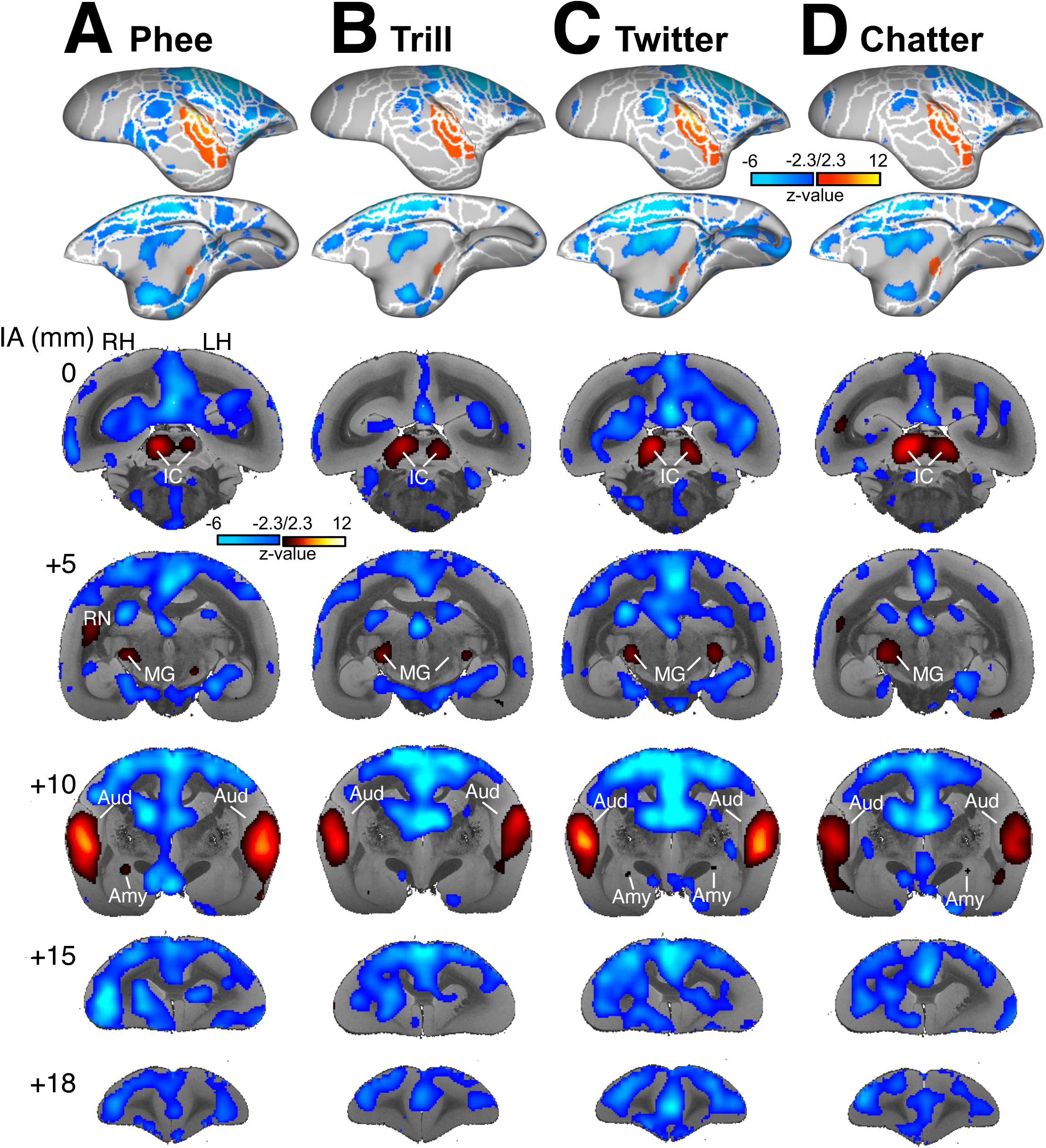
Brain activations for the different conspecific calls versus baseline. Group functional topologies (n = 9 marmosets) for Phee (A), Trill (B), Twitter (C), and Chatter (D) against baseline are presented on the lateral and medial views of the right fiducial marmoset cortical surfaces. Volumetric activations at various interaural (IA) levels are superimposed onto coronal slices of anatomical MR images. All activation maps are thresholded within the range of z-scores below −2.3 and above 2.3. Cool color gradients with negative values denote neural deactivation, whereas warm color gradients with positive values represent neural activation, illustrating both the spatial distribution and intensity of neural responses elicited by the auditory task. **LH**, left hemisphere; **RH**, right hemisphere; **Aud**, auditory cortex; **MG**, medial geniculate nucleus; **IC**, inferior colliculus; **Amy**, amygdala; **RN**, reticular nucleus of the thalamus.

At the subcortical level, the IC and MGN were activated by each of the four call types relative to the baseline. Amygdala activations were found for the presentation of phee, twitter, and chatter calls whereas the comparison between trill and baseline failed to show significant activation in this region.

To better illustrate the activations for the four calls, we displayed them on flat maps of the right hemisphere in Figure 4A. The figure shows that a few other cortical areas such as the temporo-parietal-occipital area (TPO), insular proisocortex (Ipro), and temporal proisocortex (Tpro) were activated in addition to auditory cortices by conspecific vocalizations. Moreover, cingulate areas 32, 24a-d, and 23a-c, primary motor cortex 4ab, premotor areas 6DC, 6M, 6V, 6DR, somatosensory areas 3a-b and 1/2, parietal area PE, as well as frontal area 8aD, 8Av, 8b were deactivated by presentation of each of these calls.

**Figure 4.**
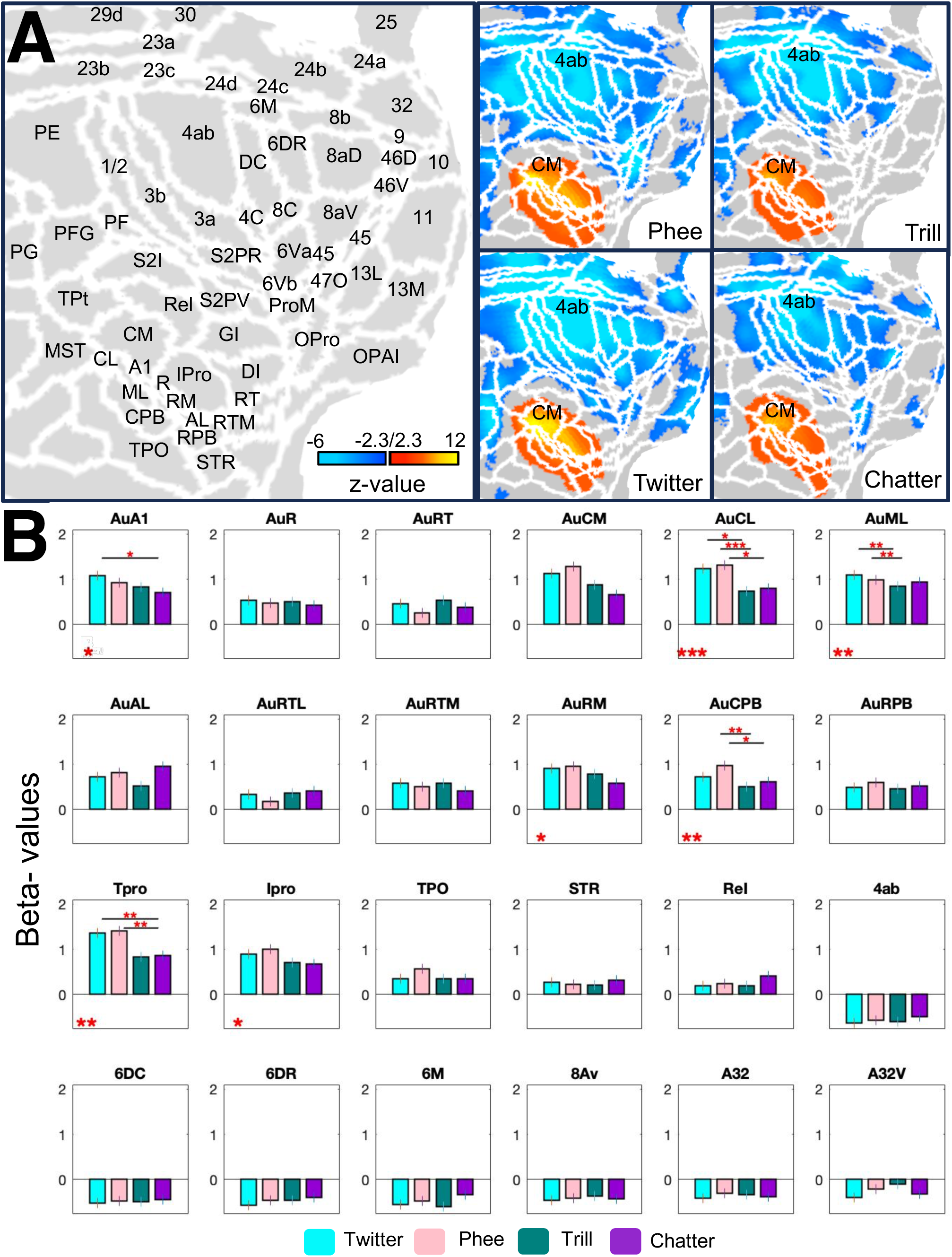
ROI analysis. **(A)** The representation of neural activity patterns for each call versus baseline on flat maps for the right hemisphere. The z-score maps are thresholded at below −2.3 and above 2.3. Warm color gradients indicate activation, while cold color gradients signify deactivation. **(B)** Beta-values analysis for the activity of each call versus baseline across 24 ROIs. Each ROI is represented by four bars related to each call, wherein the bar height reflects the magnitude of activity for the corresponding ROI in response to that specific call. Significance levels of group differences in each ROI are displayed below each graph using asterisks (* p ≤ 0.05, ** p < 0.01, and *** p < 0.001), displaying regions where differences reach statistical significance. In regions where differences between conditions are significant, asterisks indicating significance levels are displayed above the corresponding bars. (* p < 0.05, ** p < 0.01, and *** p < 0.001). The vertical line on each bar demonstrates the standard error of the mean (SEM). **CM**, auditory cortex caudomedial area; **A1**, auditory cortex primary area; **R**, auditory cortex rostral area; **RT**, auditory cortex rostrotemporal area; **CL**, auditory cortex caudolateral area; **ML**, auditory cortex middle lateral area; **AL**, auditory cortex anterolateral area; **RTL**, auditory cortex rostrotemporal lateral area; **RTM**, auditory cortex rostrotemporal medial area; **RM**, auditory cortex rostromedial area; **CPB**, auditory cortex caudal parabelt area; **RPB**, auditory cortex rostral parabelt area; **TPO**, temporo-parietal-occipital area; **Ipro**, insular proisocortex; **Tpro**, temporal proisocortex; **STR**, superior rostral temporal area; **ReI**, retroinsular area.

Despite these similarities, Fig. 4A also shows some differences in activation between the four calls. To characterize the variations in response magnitude within the call-specific network across distinct vocalizations compared to the baseline, we extracted the beta-value of each condition (i.e., phee, twitter, trill, and chatter) for 24 different regions including the auditory cortices, adjacent areas such as superior rostral temporal area (STR), TPO, Ipro, Tpro as well as primary motor cortex 4ab, premotor areas 6DC, 6M, 6V, 6DR, and cingulate areas 32 and 32V as shown in Figure 4B. Our findings highlight that despite the presence of a shared network, the magnitude of the response evoked in different parts of this network varies depending on the type of call. These findings indicate significant similarities between twitter and phee calls, showcasing a response pattern more akin to each other compared to trill and chatter calls, which exhibit a closer resemblance to one another. The bar graphs of Fig. 4B illustrate that twitter and phee calls triggered higher activation in Tpro, CL, and CM compared to other regions, while twitter also activated ML and A1 with slight variations relative to CM. Trill calls activated CM and ML, while chatter activated AL and ML more prominently compared to other regions. Our results also suggest that twitter and phee calls exhibited a more similar pattern in activation compared to trill and chatter calls, which were more alike.

To better understand the group and condition differences, we conducted repeated measures analysis of variance (rmANOVA). Significant within-group distinctions across several regions, including CL, ML, CPB, Tpro, RM, A1, and Ipro were identified. The differences with a significant p-value were displayed by asterisks beneath each region of interest (ROI) in Figure 4B. We found significant differences between the calls in CL (F_(3,243)_ = 6.66, p < 0.001), ML, CPB, Tpro (F_(3,243)_ = 4.95, F_(3,243)_ = 5.13, F_(3,243)_ = 5.56, respectively, all p < 0.001), as well as in RM, Ipro, and A1 (F_(3,243)_ = 2.82 and F_(3,243)_ = 3.02, F_(3,243)_ = 2.63, respectively, all p ≤ 0.05). Between-condition differences are indicated above corresponding bars in each of these ROIs, where significant differences were found between twitter and chatter in A1 (p < 0.05) and Tpro (p < 0.01), between twitter and trill in CL (p < 0.05) and ML (p < 0.01), between phee and trill in CL (p < 0.001), ML (p < 0.01), and CPB (p < 0.01), and between phee and chatter in CL (p < 0.05), CPB (p < 0.05), and Tpro (p < 0.01).

Within the auditory cortex, twitter calls induced stronger activations in belt areas such as ML, CM, CL, and RM compared to other regions. For phee calls, the most substantial activation in the auditory cortex was observed in belt areas including CL, CM, ML, RM, and the caudal parabelt (CPB). The response of different regions within the auditory cortex was almost identical for trill and chatter calls, with the highest level of activation observed in CM and AL for each respective call, along with ML showing significant activation for both types of calls. Additionally, RTL in the auditory cortex consistently exhibited the lowest level of activation across twitter, phee, and trill stimuli, whereas chatter calls evoked the lowest activation in the RT field. Adjacent areas such as TPO, Ipro, Tpro, STR, and the retroinsular area (ReI) also showed variations in their activation levels. In addition, all calls deactivated primary motor cortex 4ab, premotor areas 6DC, 6M, 6DR, frontal area 8Av, and cingulate areas 32 and 32V. The greatest deactivation across these regions was observed in area 4ab for all calls. Moreover, area 32 experienced the highest degree of deactivation in response to twitter, while other calls induced relatively similar levels of deactivation in this region (See Supplementary Figure 2).

To identify call-specific activations, we compared the responses to each call with those of the other three calls. Figure 5 depicts these results both on flat maps and in volume space (coronal and sagittal views). Phee calls (1^st^ row) were characterized by more robust activations in the auditory area CPB and area TPO. In the cerebellum, phee calls elicited stronger responses in the deep cerebellar nuclei (DCN) and cerebellar lobules VIIB and VIIIA. For trill calls (2^nd^ row), larger responses were present in orbitofrontal area 11 and anterior cingulate cortex area 32 compared to the other calls. Twitter calls (3^rd^ row) exhibited stronger activation in area CM of the auditory cortex and the MG. Finally, chatter calls (4^th^ row), were associated with more intense activations in the superior colliculus (SC), parietal area PE, and premotor area 6M in comparison to the other vocalizations.

**Figure 5.**
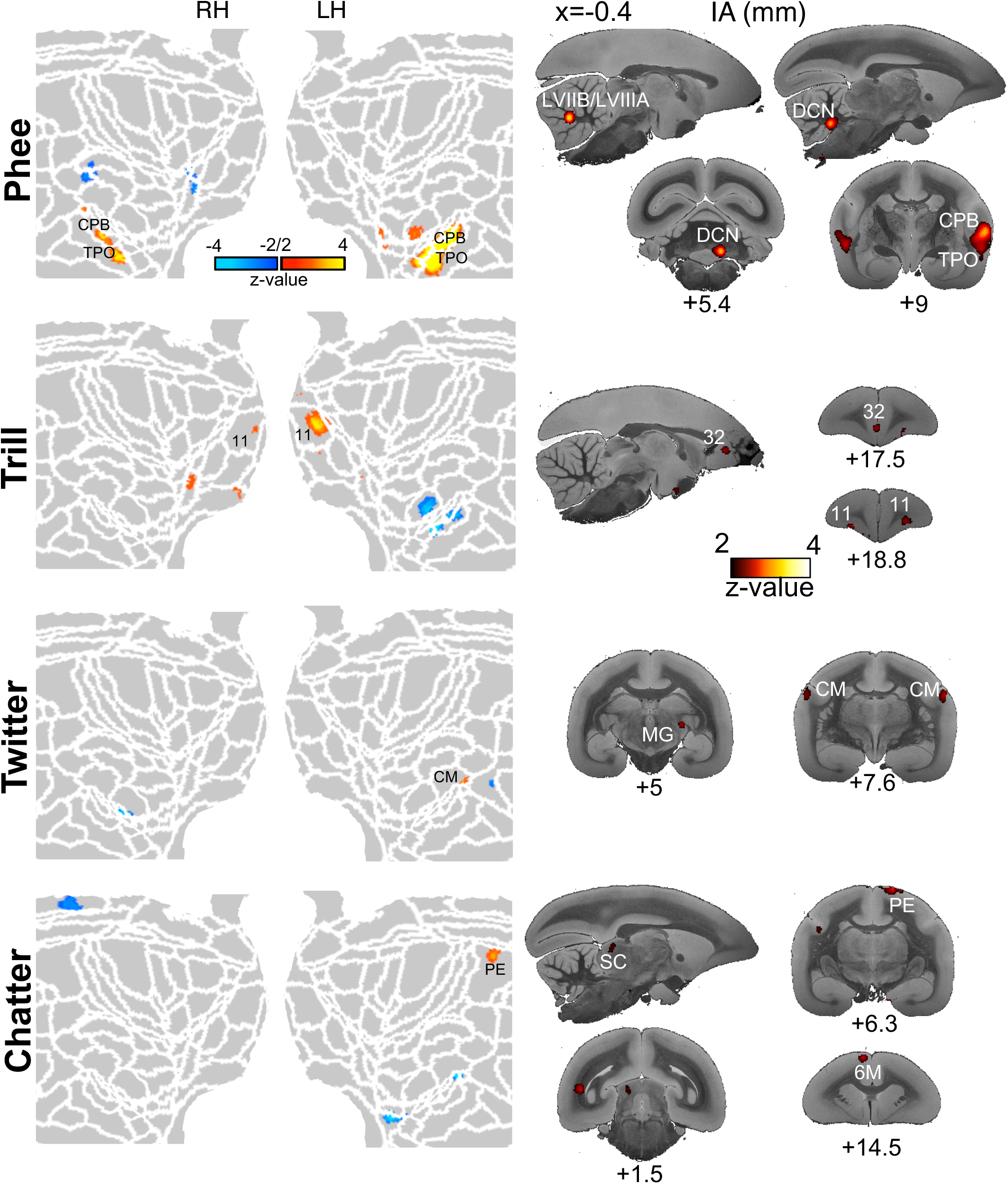
Brain activity in response to each call versus all other calls. Each row illustrates the neural response of brain regions in response to phee, trill, twitter, and chatter calls, respectively, compared to all other calls. The neural patterns are displayed on flat cortical maps for both the right and left hemispheres, along with volumetric representations at different interaural (IA) levels. All z-score maps are thresholded to display values < −2 and > 2. **LH**, left hemisphere; **RH**, right hemisphere; **CPB**, auditory cortex caudal parabelt area; **TPO**, temporo-parietal-occipital area; **DCN**, deep cerebellar nuclei; **LVIIB**/**LVIIIA**, cerebellar lobules VIIB and VIIIA; **CM**, auditory cortex caudomedial area; **MG**, medial geniculate nucleus; **SC**, superior colliculus; **PE**, parietal area.

We previously showed that blocks of vocalizations compared to scrambled vocalizations or nonvocal sounds activate the medial prefrontal cortex (mPFC) area 32. Therefore, we compared activations in mPFC between the calls (Fig. 6). The calls on the x-axis were compared to those listed on the y-axis (calls _x-axis_ > calls _y-axis_), and the results were thresholded at z-scores higher than 2. The results show that area 32 was significantly more activated by phee and trill calls than twitter calls. Area 9 was more activated by trill and phee calls than by chatter calls and area 14R was more active for chatter than phee or twitter calls. Overall, the results show more activation in the mPFC cortex for phee and trill calls compared to other calls.

**Figure 6.**
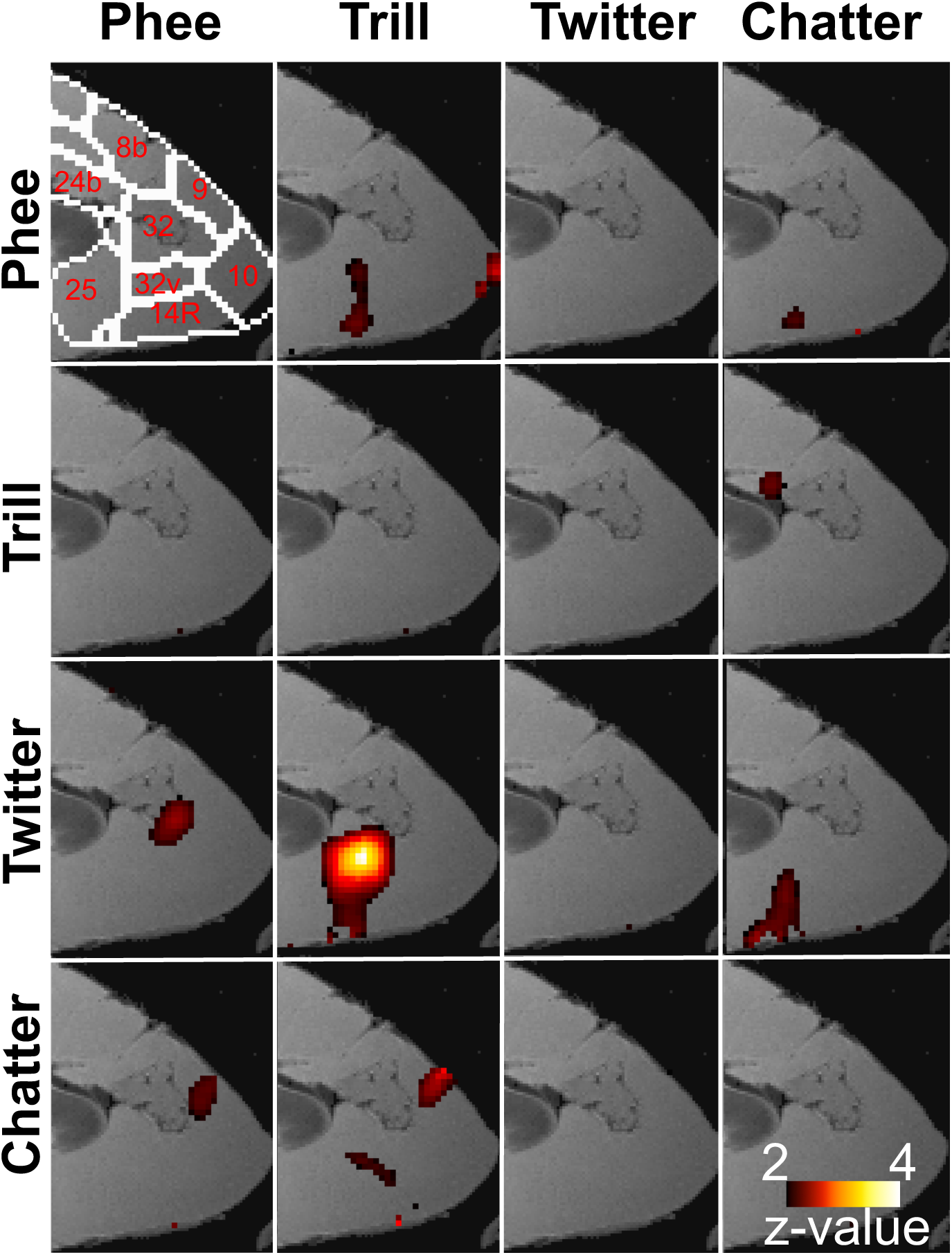
Neural response of medial prefrontal cortex (mPFC) in response to each call versus each other calls. The volumetric representation of the neural activity of mPFC for calls on the x-axis compared to those listed on the y-axis (calls _x-axis_ > calls _y-axis_). Results are thresholded at z-scores higher than 2.

## Discussion

Our recent research revealed that conspecific vocalizations activate a fronto-temporal network in marmosets^15^. In the present study, we investigated the specificity of this network in processing various calls by conducting whole-brain event-related fMRI in nine adult marmosets. The sounds encompassed four prevalent call types: phee, chatter, twitter, and trill. Consistent with our prior findings ^15^ and existing studies on the marmoset auditory pathway ^16^, we found that the calls activated a network, encompassing both cortical and subcortical structures. The cortical regions involved primarily auditory cortices, including its core, belt, and parabelt areas, as well as adjacent areas like TPO, Ipro, Tpro, STR, and ReI.

Previous studies have examined the amygdala’s role in processing auditory stimuli ^17,18^. It has been shown that this processing is dependent on the context of the presented sounds, where a specific sound can either activate or deactivate the amygdala ^19^. If the activation of the amygdala is related to the acoustic features, it would be expected that the amygdala responded to both trill and phee calls in our experiment, given their similar acoustic features including spectral, temporal, and amplitude characteristics ^20,21^. However, our observations diverged from this expectation. We found that while exposure to phee, twitter, and chatter calls specifically activated the amygdala, trill calls did not induce notable activation in this region, supporting the notion of the amygdala’s context-dependency function.

When comparing all vocalizations against the baseline, as illustrated in Figs. 1C and 2, we observed that many cortical regions, including premotor cortical areas 6DC, 6DR, 6Va, 6M, primary motor cortex 4ab, cingulate cortices 24a-d, 32, 23a-c, 23v, prefrontal area 8Av, and somatosensory cortex areas 1/2 and 3, were deactivated in response to short conspecific calls. These areas are known for their roles in controlling motor functions including vocalization, such as the voluntary initiation of vocalizations in non-human primates (NHPs) and speech in humans ^22^ ^23^.

Additionally, our results, as shown in Fig. 2, show deactivation of the cerebrocerebellum which is known to play an important role in motor preparation ^24^. These deactivations suggest that the marmosets likely reduced minor limb and body movements present during the baseline (lasting 3-12 seconds) when exposed to the shorter vocalizations (0.6 seconds). It is possible that the animals entered a state of stillness, suppressing motor activity to focus on perceiving the auditory stimuli. This finding aligns with a functional ultrasound study that reported reduced blood flow in the medial sensorimotor cortex (mSMC) during the presentation of conspecific vocalization in marmosets ^8^.

Several functional magnetic resonance imaging (fMRI) experiments with awake marmosets have revealed a tonotopic organization in their auditory cortex. This tonotopic arrangement is consistent with the general pattern observed in the auditory cortex of other primates, corroborating findings from optical recordings ^25^ and electrophysiological studies ^26^. When exposed to pure tones and bandpass-filtered noise containing both low and high frequencies, high frequencies (4 – 16 kHz) activated the caudal region in area A1 and the border of areas R and RT, whereas low frequencies (0.25 – 1 kHz) activated at the border of A1 and R, as well as the rostral end of RT ^27^. Our data, illustrated in Figure 4A, is in line with this tonotopic structure. We identified a caudal-rostral gradient for vocalization selectivity in the auditory cortex. Calls with high frequencies, such as twitter, phee, and trill, produced stronger activation in the caudal portion of the primary auditory cortex A1 and the caudal regions of the belt field such as CL and CM. In contrast, the low-frequency chatter call elicited greater activation in AL and ML. (See Supplementary Figure 2)

Furthermore, our research indicates that long-distance calls such as twitter and phee engage more similar brain regions. Moreover, narrow-band calls such as trill and phee, often grouped together due to their acoustic similarities, showed less similarity in neural responses. This observation shows that the functional activity of brain regions relies significantly on the context of the calls, rather than solely on their acoustic features. This finding is also in line with a previous study using functional ultrasound imaging which did not find statistically significant differences in cerebral blood volume rates in medial brain regions for twitter and phee calls. However, trill calls and alarm calls exhibited significant differences, particularly in the mPFC ^8^.

The presence of call-specific selectivity in CL, ML, and AL regions of the macaque auditory cortex ^2,4^ prompted the question of whether a similar hierarchical organization exists in marmosets. We found notable group distinctions in CL (p < 0.001), ML and CPB (both p < 0.01), RM (p < 0.05), and A1 (p = 0.05) within the auditory cortex, indicating discrimination of conspecific calls in these regions, as depicted in Fig. 4B.

Our findings also suggest that the cerebellum plays a particularly important role in processing phee calls. Unlike other vocalizations exchanged in direct contact or close proximity among marmosets, phee calls are communicated over long distances or between individuals occluded from visual contact. Also, previous studies suggest a diverse acoustic attribute associated with different types of phee calls, and their categorization hinges on factors such as the physical distance between the callers and their identity ^28^. More demanding processing needs for phee calls are also supported by our observation that the caudal parabelt (CPB) and area TPO responded higher to phee calls than to other calls.

A prominent activation was seen in the anterior cingulate area 32 and medial orbitofrontal cortex 11 (mOFC 11) for trill calls in comparison with all other conditions in our results. Trill calls are typically exchanged at a close distance between social partners during relaxed social states like foraging and resting and are usually regarded as positive welfare indicators ^9,29,30^. Indeed, medial orbitofrontal cortex area 11 is known for its responsiveness to pleasant stimuli in macaques and it is connected to the pregenual anterior cingulate cortex ^31^.

In our study, twitter calls elicited increased bilateral activation in CM and more pronounced activity in the left hemisphere of MG than in other conditions. Recent research proposes that CM neurons are integral to temporal processing, especially in the precise discrimination of temporal envelopes ^32^. Intriguingly, psychophysical experiments involving human subjects and speech stimuli have indicated that features of the temporal envelope modulation, rather than those of the spectral envelope, are crucial in speech identification and recognition, both in quiet and noisy environments ^33^. Given that twitter calls are agonistic intergroup calls ^34^, their activation of the CM suggests that discrimination of the temporal envelope may play a key role in sound identification.

For chatter calls, our observations showed significant activity in the SC, parietal area PE, and premotor area 6M, more than for other conditions. This pattern of neural activity suggests a specific engagement of these areas in processing the unique characteristics of chatter calls, potentially indicating responsive orienting reactions to such calls.

Previously, we found strong activations for longer periods of vocalizations (12 sec) in mPFC area 32. Although we show here that all brief call types (0.6 sec) suppressed activations in the mPFC in comparison to the long baseline periods (3 - 12 sec), we found significantly higher activity in response to phee and trill calls when compared with twitter and chatter calls. These two calls belong to the category of communicative calls, often grouped together ^20^. This finding points to an important role in area 32 for these communicative calls which are used when marmosets engage in antiphonal calling.

In conclusion, our event-related fMRI study has revealed several key aspects of vocalization processing in marmosets. In addition to the finding of a core cortical and subcortical network activated by phee, trill, twitter, and chatter vocalizations, the observed deactivations in certain cortical and subcortical areas during vocalization aligns with the notion of reduced motor activity to facilitate enhanced auditory perception. In addition, the cerebellum’s significant role in processing phee calls, the activation patterns in the anterior cingulate and orbitofrontal cortices for trill calls, and the involvement of the caudomedial belt and medial geniculate in processing twitter calls all point to a highly specialized and context-dependent vocalization processing system in marmosets.

## Materials and Methods

### Animal subjects

All experimental procedures complied with the guidelines outlined by the Canadian Council of Animal Care and were conducted in accordance with a protocol adhered by the Animal Care Committee of the University of Western Ontario. In this study, we conducted whole-brain fMRI scans on nine adult common marmosets (*Callithrix jacchus*: four females, two left-handed, age: 48 - 74 months, and weights: 380 - 464 grams). All animals were housed in pairs in a colony located at the University of Western Ontario where environmental conditions included a humidity level of 40 - 70%, a diurnal 12-hour light cycle, and a temperature maintained between 24 - 26° C.

### Surgical procedure

Animals underwent an aseptic surgery for implanting a compatible machined PEEK (polyetheretherketone) under anesthesia to prevent any head motion while scanning. During the surgical procedure, the marmosets were sedated and intubated to be maintained under gas anesthesia (a mixture of O_2_ and air, with isoflurane levels ranging from 0.5% to 3%). The skull surface was initially prepared by applying several coats of adhesive resin (All-Bond Universal; Bisco, Schaumburg, Illinois, USA) after a midline incision of the skin along the skull. The resin-coated area was air-dried and then cured with an ultraviolet dental curing light (King Dental). Then the PEEK head post was secured on the skull using a resin composite (Core-Flo DC Lite; Bisco, Schaumburg, Illinois, USA). Throughout the surgery, continuous monitoring was conducted to track heart rate, blood pressure, oxygen saturation, and body temperature. More details regarding the animal surgery protocol are included in Johnston *et al.*, 2018 ^35^ and Zanini *et al.*, 2023 ^36^.

### Animal habituation

Following a recovery period of two weeks after the surgery, the monkeys were gradually acclimatized to the head-fixation system and the MRI environment within a mock scanner. Specific details concerning our training protocol are found in Zanini *et al.*, 2023 ^36^.

### Marmoset contact calls and their characteristics

Previous studies in marmosets, whether in captivity or freely moving, indicate that their vocal repertoire comprises ∼13 distinct call types, each capable of eliciting varied behavioral responses from conspecific listeners ^9^. These vocalizations are influenced by social contexts and can serve as indicators of their overall welfare ^8,9,37^. Different acoustic factors such as bandwidth, harmonic ratio, and duration vary among individual marmosets and even within calls of the same animal for a specific call type ^38^. A fundamental aspect of exchanging calls among marmosets is the temporal adjustment of contact calls, also known as turn-taking. Within the array of marmoset vocalizations, the trill and phee calls serve the purpose of antiphonal communication and are considered a key factor in facilitating the turn-taking aspect ^10^. Trill calls, as a short-distance call, are the most dominant vocalizations within the marmoset repertoire. They are produced when individuals are in close vicinity to each other. Previous vocal interaction studies demonstrated that high-frequency trill calls predominantly occur between partners and are less frequently exchanged with other members within the group. Additionally, trills emitted by one animal are often followed by trills from other animals within a period of less than one second. Conversely, high-frequency phee calls are emitted when a marmoset is distant from its peers, lacking visual contact ^9–11^. Phee calls can be different in their acoustic features and are classified based on the distance of the callers, whether it is longer or shorter, from their conspecifics ^28,39^. Twitter calls are characterized as loud sounds that share a structural resemblance with warbles. These high-frequency calls consist of a sequence of several brief, rapidly frequency-modulated call phrases with relatively consistent inter-syllable intervals, typically produced during intergroup agonistic interactions ^9,11,13,34,40^. Unlike phee calls, twitter calls are classified as short-distance calls due to their low amplitude, which might not effectively transmit over long distances ^41^. Low-frequency chatter calls are also produced by marmosets, serving as a manifestation of distinct emotional contexts including intergroup or outgroup aggression ^39^.

### Stimuli generation

The natural vocalizations used in this study were recorded in a small colony that accommodated five groups of marmosets under this study or their companions. Monkeys were paired-housed in five different cages (2 - 6 individuals/cage) at the University of Western Ontario. Conspecific vocalizations frequently produced in the vocal exchange between members of the colony were recorded over a period of two hours using a microphone (MKH 805, Sennheiser, Germany in combination with phantom power, NW-100, NEEWER) connected to a laptop (Macbook Pro, Apple) and stored as Wave files in Audacity software (v3.2.5) ^42^. No manipulation such as filtering or background noise removal was performed on the archived data to preserve the integrity of vocal features. Then, data were explored through its spectrogram using the Audacity software package ^42^, leading to the identification of three distinct categories of social calls including phee, twitter, and trill calls. Among the recorded data, twitter and trill were the most prevalent calls. Instances of calls with low amplitude or intensity, which likely originated from animals housed in distant cages, were excluded from our selected dataset. For trill and twitter calls, we selected two samples each from our recorded calls. However, we opted to use other sources and pre-recorded samples for phee and chatter (n = 2) calls ^10^ due to their suboptimal quality, primarily resulting from overlapping individual calls with background noise or vocalizations from multiple monkeys, or the absence in our original recordings. The spectral power of candidate samples was subsequently normalized using a custom MATLAB script and was matched in duration for 600 ms using online AudioTrimmer. Ultimately, we employed a collective count of eight samples of the four mentioned call categories (two samples from each of trill, chatter, twitter, and phee calls) as our vocal stimuli in this study. The spectrograms of all auditory stimuli, which were employed in the current study, are displayed in Supplementary Figure 1.

### fMRI data parameters

Each fMRI session lasted 40 - 55 minutes depending on whether marmosets were scanned for two or three functional runs using gradient-echo-based single-shot echo-planar images (EPI) sequences. To minimize the impact of the noise emanating from the scanner on auditory stimuli, we adopted a “bunched” acquisition approach. In this method, the acquisition of image slices occurred within a period (*TA*) that was shorter than the repetition time (*TR*), thus leaving a silent interval (*TS*) during which the auditory stimuli were presented to the animals (*TR = TA+TS*). The parameters used for data collection in this experiment are as follows: repetition time (*TR*) = 3 s, acquisition time (*TA*) = 1.5 s, silent period (*TS*) = 1.5 s, *TE* = 15 ms, flip angle = 40°, field of view = 64×48 mm, matrix size = 96×128, voxel size = 0.5 mm isotropic, number of axial slices = 42, bandwidth = 400 kHz, GRAPPA acceleration factor (left-right) = 2. Furthermore, during each fMRI session, an additional series of EPIs was acquired with an opposing phase-encoding direction (right-left). This was done to reduce spatial distortion induced using *topup* in FSL ^43^ in our subsequent analysis. For each animal, we also collected a T2-weighted anatomical image in a separate session to facilitate the anatomical registration of the fMRI data. The T2-weighted anatomical image was obtained using the following imaging parameters: *TR* = 7 s, *TE* = 52 ms, field of view = 51.2×51.2 mm, voxel size = 0.133×0.133×0.5 mm, number of axial slices = 45, bandwidth = 50 kHz, GRAPPA acceleration factor = 2 ^36^.

### fMRI setup

Functional imaging data was acquired at the Centre for Functional and Metabolic Mapping (CFMM) at the University of Western Ontario, using an ultrahigh-field magnetic resonance system operating at 9.4 Tesla, featuring a 31 cm horizontal bore magnet and a Bruker BioSpec Avance III console running the Paravision 7 software package. A custom-designed gradient coil ^44^ with an inner diameter of 15 cm, boasting a maximum gradient strength of 400 mT/m, was employed. Additionally, an 8-channel receive coil ^45^ was positioned inside a home-built transmit quadrature birdcage coil with an inner diameter of 120 mm. A comprehensive description of the animal’s preparation can be found in ^15,36^. Vocal stimuli were presented passively during each run using a custom-built MATLAB script (R2021b, The MathWork Inc.). A transistor-transistor logic (TTL) pulse box was used to synchronize the onset of the echo-planar imaging (EPI) sequences with the onset of the auditory stimuli and video recordings.

### fMRI task design and sound presentation

The fMRI task involved passively presenting the marmosets with the vocal stimuli described above. During the sessions, each marmoset was scanned for a total number of 2 - 3 runs. Each run contained 300 functional volumes (*TR* = 3 s). This resulted in a total of 86 functional runs of which 4 runs were excluded and not incorporated into our final dataset due to technical issues during scanning sessions where vocal stimuli were not properly presented.

To present vocal stimuli, a customized MATLAB script in line with the event-related imaging paradigm was employed. This script controlled the initiation of sound presentation through a TTL pulse received from the MR scanner. Each scanning session started and ended with a 1.5-second baseline period during which no auditory stimuli were introduced. Using a silent period (*TS*) of 1.5 seconds within the *TR* enabled us to present stimuli without interference from scanner noise. Subsequently, a specific call was chosen in a random order and played back during *TS*, with the interstimulus intervals being selected in a pseudorandom sequence from a predefined set of intervals including 3, 6, 9, and 12 seconds.

## Data Analysis

### Preprocessing of fMRI data

The image preprocessing for this study involved the application of a general linear model analysis through AFNI ^46^ and FSL ^43^. Prior to initiating the preprocessing of the functional data, the T2-weighted anatomical data were reoriented. Subsequently, a manual skull-stripped mask was created for each individual subject using FSLeyes application ^43^ and then converted into a binary mask. This binary mask was then multiplied with the anatomical data to produce the T2 template mask and ultimately registered to the 3D NIH marmoset brain atlas (NIH-MBA) ^47^ through Advanced Normalization Tools (ANT’s *ApplyTransforms*) ^48^ using a non-linear registration. Functional data preprocessing briefly includes: 1. Conversion of raw functional data from DICOM to NIfTI formats (*dcm2niix*) 2. Reorientation of the functional data (*fslswapdim)* 3. Registering functional images to anatomical data (*fslroi and topup*) 4. Interpolating each run (*applytopup*) 5. Removal of spikes (*3dToutcount* and *3dDespike*) 6. Time shifting (*3dTshift*) 7. Registration of the entire dataset to a reference volume, typically the middle volume (*3dvolreg*) 8. Spatial smoothing by convolving the BOLD image with a three-dimensional Gaussian function having a full-width-half-maximum (FWHM) of 1.5 mm (*3dmerge*) 9. Bandpass filtering within the frequency range of 0.01 to 0.1 Hz (*3dbandpass*) 10. Calculation of a mean functional image for each run which was linearly aligned with the corresponding T2-weighted anatomical image of each animal (FSL’s *FLIRT*). The resulting transformation matrix from the realignment process was then employed to transform the 4D time series data.

### Statistical analysis of fMRI data

To generate the General Linear Model, the event onsets that were already obtained during scans were convolved to the hemodynamic responses (AFNI’s Convolution*, GAM(4,0.7)*). A regression was then generated for each condition which was used for regression analysis (*3dDconvolve*) along with polynomial detrending regressors and the motion parameters acquired from previous preprocessing steps. The resultant regression coefficient maps were then registered to template space using the transformation matrices obtained with the registration of anatomical images on the template. Then, one T-value map for each call per run was obtained after registering to the NIH Marmoset Brain Atlas ^47^. The obtained maps were subsequently subjected to group-level comparisons through paired t-tests (*3dttest++),* yielding Z-value maps. These functional maps were then visualized on fiducial maps using Connectome Workbench v1.5.0 ^49^ as well as coronal and sagittal sections using FSLeyes ^43^. To determine the locations of activated brain regions, we aligned these functional maps with a high-resolution (100 × 100 × 100 μm) ex-vivo marmoset brain ^50^, registered on the NIH marmoset brain template ^47^, and employed Paxinos parcellation for region identification ^51^.

### Quantification of local responses

To evaluate how the brain responds to different types of vocalizations, we analyzed the changes in neural activation across 24 ROIs, including auditory areas and peripheral regions. We extracted time-course data from these ROIs using AFNI’s *3dmaskave* tool for each of the 82 experimental runs across nine monkeys. Next, we calculated the average response and standard error of the mean (SEM) across all runs for each ROI using a custom MATLAB script. Additionally, we conducted a repeated measures analysis of variance (rmANOVA) on the time-course data of each region in MATLAB to examine differences between experimental conditions (*ranova, multcompare*). This analysis helped us determine the significance of observed neural activity patterns ^52^.

## Acknowledgments

We wish to thank Cheryl Vander Tuin, Whitney Froese, Miranda Bellyou, and Hannah Pettypiece for animal preparation and care and Dr. Alex Li for scanning assistance. Support was provided by the Canadian Institutes of Health Research (<u>FRN 148365</u>, S.E.), the Natural Sciences and Engineering Council of Canada (S.E.), and the Canada First Research Excellence Fund to BrainsCAN. We also acknowledge the support of the Government of Canada’s New Frontiers in Research Fund (NFRF), [NFRF-T-2022-00051].

