## Supplementary figures for "Unique cortical and subcortical activation patterns for different conspecific calls in marmosets"

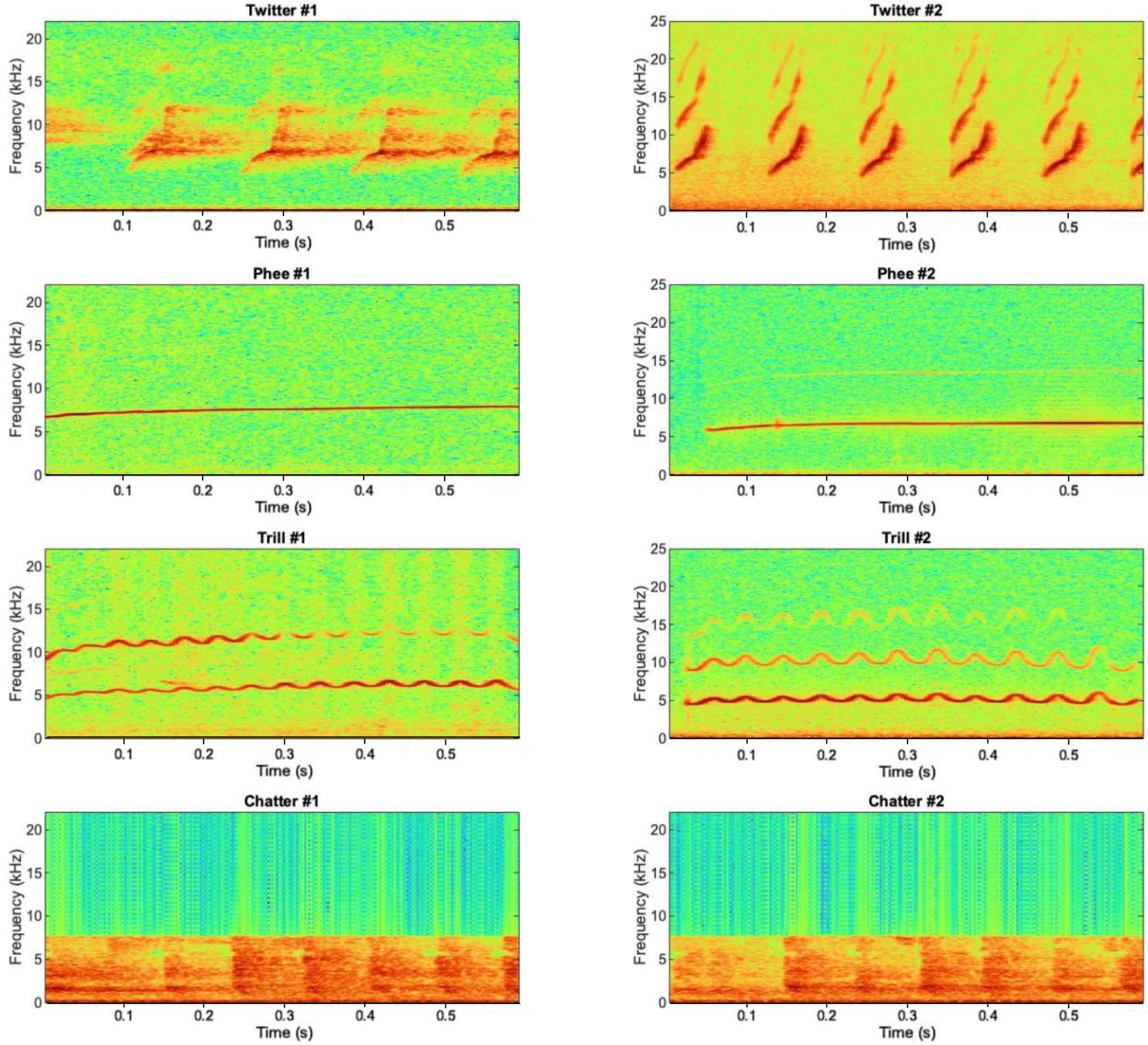


**Supplementary Figure 1. The spectrogram of all vocal stimuli employed for this study.** The length of all vocal stimuli was set to 0.6 seconds. Vocalizations such as twitter, phee, and trill are classified as high-frequency, while chatter falls into the category of low-frequency calls.

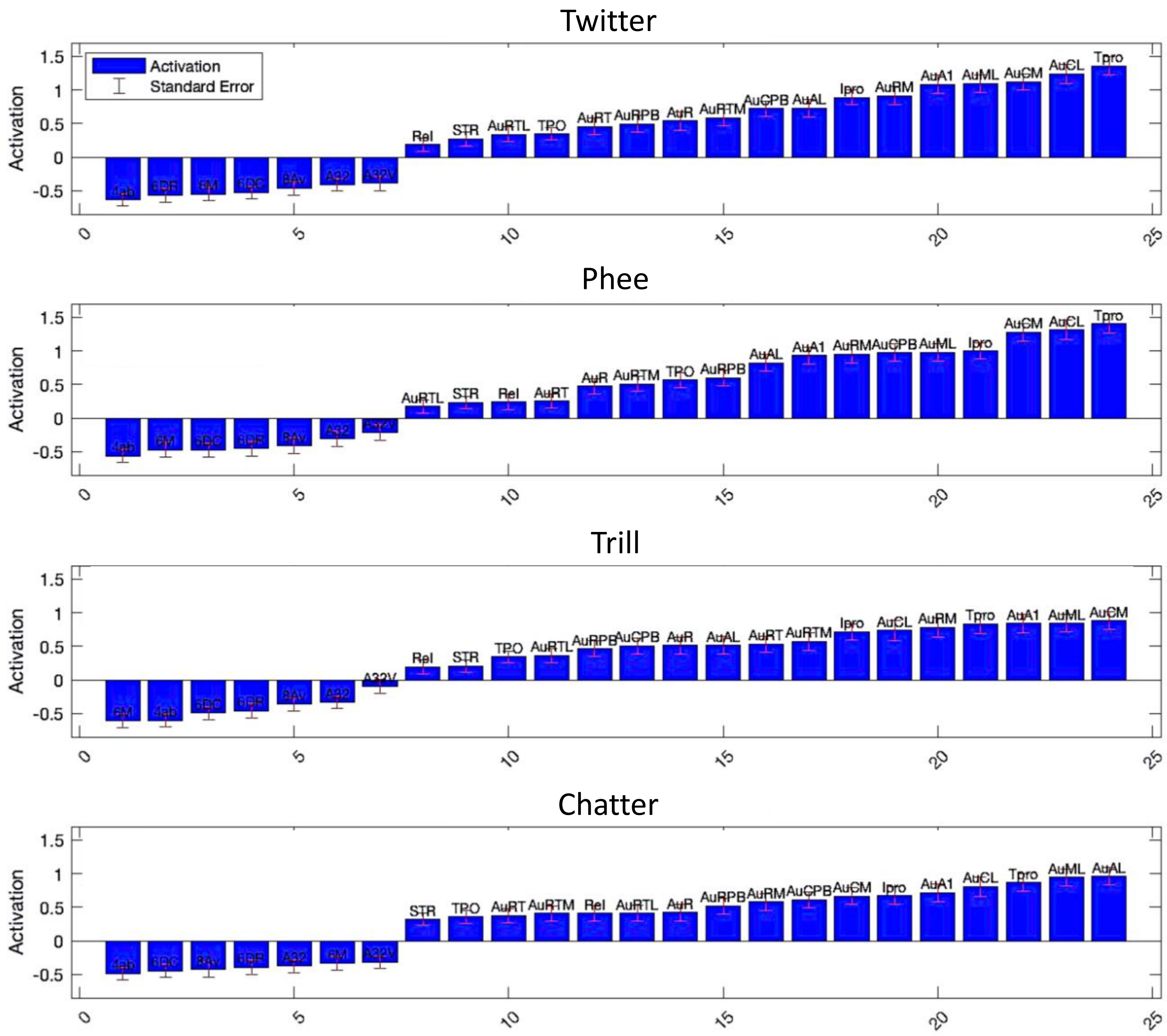


**Supplementary Figure 2. The order of activity strength in different ROIs for each call.** The figure illustrates the order of activity strength for each ROI across each call. Each ROI is represented by a bar graph, wherein the height of the bars reflects the magnitude of activity. The red vertical line on each bar demonstrates the standard error of the mean (SEM). **CM**, auditory cortex caudomedial area; **A1**, auditory cortex primary area; **R**, auditory cortex rostral area; **RT**, auditory cortex rostrotemporal area; **CL**, auditory cortex caudolateral area; **ML**, auditory cortex middle lateral area; **AL**, auditory cortex anterolateral area; **RTL**, auditory cortex rostrotemporal lateral area; **RTM**, auditory cortex rostrotemporal medial area; **RM**, auditory cortex rostromedial area; **CPB**, auditory cortex caudal parabelt area; **RPB**, auditory cortex rostral parabelt area; **TPO**, temporo-parietal-occipital area; **Ipro**, insular proisocortex; **Tpro**, temporal proisocortex; **STR**, superior rostral temporal area; **ReI**, retroinsular area.
